# Large-scale determination and characterization of cell type-specific regulatory elements in the human genome

**DOI:** 10.1101/176602

**Authors:** Can Wang, Shihua Zhang

**Affiliations:** National Center for Mathematics and Interdisciplinary Sciences, Academy of Mathematics and Systems Science, Chinese Academy of Sciences, Beijing 100190, China; School of Mathematics Sciences, University of Chinese Academy of Sciences, Beijing 100049, China

## Abstract

Histone modifications have been widely elucidated to play vital roles in gene regulation and cell identity. The Roadmap Epigenomics Consortium generated a reference catalogue of several key histone modifications across >100s of human cell types and tissues. Decoding these epigenomes into functional regulatory elements is a challenging task in computational biology. To this end, we adopted a differential chromatin modification analysis framework to comprehensively determine and characterize cell type-specific regulatory elements (CSREs) and their histone modification codes in the human epigenomes of five histone modifications across 127 tissues or cell types. The CSREs show significant relevance with cell type-specific biological functions and diseases and cell identity. Clustering of CSREs with their specificity signals reveals diverse histone codes, demonstrating the diversity of functional roles of CSREs within the same cell or tissue. Last but not least, dynamics of CSREs from close cell types or tissues can give a detailed view of developmental processes such as normal tissue development and cancer occurrence.

## Introduction

Different cells in human behave distinctly and perform diverse functions. However, DNA sequences within nucleus are the same. Histones bound by DNA can carry various post-translational modifications, most of which have high plasticity across cell types on different locations of genome, giving an additional layer of information to DNA sequence (Mikkelsen et al., 2007). A plenty of histone modifications help chromatin to encode various programs of gene regulation, resulting in different protein functions and cell type-specific phenotypes. A specific histone modification influences chromatin compaction and accessibility, relating to transcription initiation and elongation, transcription factor binding, enhancer activation and repression, etc (Lawrence et al., 2016).

By leveraging next generation sequencing, the Roadmap Epigenomics Consortium generated a reference catalogue of human epigenomes across a variety of tissues and cell types, involving a plethora of histone modifications, as well as DNA methylation, chromatin openness and RNA expression (Kundaje et al., 2015). These genome-wide maps of human epigenomic marks give rise to investigating cell type-specific functions and diseases in a more detailed manner.

Given signals of multiple histone modifications along the whole genome for a specific cell type, a variety of tools such as ChromHMM, Segway, EpiCSeg, GenoSTAN (Ernst and Kellis, 2012; Hoffman et al., 2012; Mammana and Chung, 2015; Zacher et al., 2017) have been designed to segment chromatin into different functional elements, usually called chromatin states. They can annotate multiple cells by concatenating all genomes together. However, this may miss the detailed patterns that only rarely appear in a specific cell type. Some other tools, like TreeHMM, hiHMM and IDEAS (Biesinger et al., 2013; Sohn et al., 2015; Zhang et al., 2016), give more freedom to each sample and can define chromatin states of multiple cells simultaneously. However, these tools were not ready to identify tissue- or cell type-specific regulatory elements.

Some efforts have been devoted to detect regulatory elements base on chromatin states. The Roadmap Epigenomics Project used *k*-means algorithm to clustering regulatory regions, but they only focused on enhancers and promoters, ignoring specific repressed regions. ChromDiff (Yen and Kellis, 2015) compared proportions of chromatin states along a gene between two groups to identify differential genes. However, this method cannot detect genes harboring chromatin states with the same proportion but different location distributions. EpiCompare (He and Wang, 2017) is a web tool to identify enhancers between two groups of epigenomes based on chromatin states or binary peaks of a histone modification, which must lose information among the original signals. Tools involving dPCA (Ji et al., 2013) and MultiGPS (Mahony et al., 2014) can directly compare signals between two conditions. While they cannot identity cell type-specific regions from groups of tissues or cell types.

Chen et al. (2013) proposed dCMA to identify *de novo* cell type-specific regulatory elements (CSREs) on a genome-wide scale by ChIP-seq data of eight histone modifications along with CTCF and a control across nine cell lines. Their early analysis on CSREs show that epigenomic modifications function in cell type-specific manners. However, their result would be biased by the small number of cell types analyzed. Moreover, the method was only designed for binarized data which would weaken very high signals. Large-scale determination and characterization of cell type-specific regulatory elements in the human genome is still urgently needed.

To this end, we adopted a differential chromatin modification analysis framework to comprehensively determine and characterize CSREs and their specificity signals in the human epigenomes of five histone modifications across 127 tissues or cell types. The CSREs show significant relevance with cell type-specific biological functions, diseases and cell identity. Clustering of CSREs with their specificity signals reveals diverse histone codes, demonstrating the diversity of functional roles of CSREs in the same cell or tissue. Last but not least, dynamics of CSREs from close cell types can give a detailed view of developmental processes such as normal tissue development and cancer occurrence.

## Results

### Large-scale determination of CSREs

We adopted a computational pipeline to systematically identify CSREs from the epigenomic landscape of the 127 cells or tissues in a comparative manner (Materials and methods, Supplementary Figure S1 and Table S1). We developed a web tool (http://zhanglabtools.org:2017/csre) to visualize CSREs and help users to browse their interested regions or genes to see whether they are related to CSREs or not.

In total, we identified ∼3670 CSREs per cell type or tissue on average, spanning ∼8.1 million base pairs (bp) on average (0.27% of the genome) (Figure 1A and Supplementary Table S2). The median lengths of CSREs across tissues or cell types ranging from 1625 to 2250 bp. However, the total length of CSREs varies from 38.1 Kb (small intestine) to ∼56.28 Mb (monocytes-CD14+ RO01746 primary cells). The number of CSRE neighboring transcripts (or say genes) ranges from 41 (small intestine) to 16035 (monocytes-CD14+ RO01746 primary cells) (Supplementary Figure S2). The diversity of these distributions indicates the functional complexity of human tissues or cell types.

**Figure 1.**
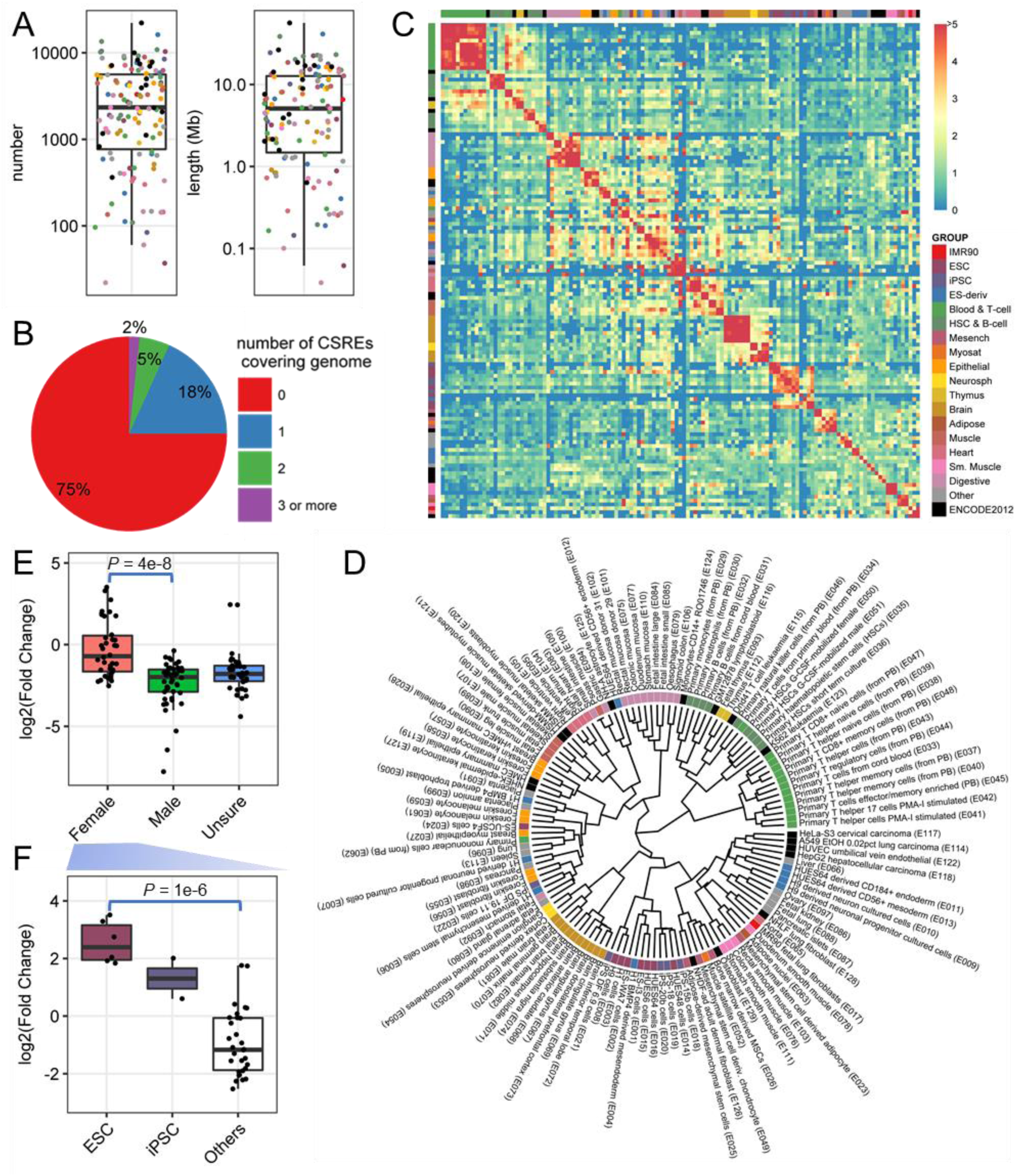
Basic characteristics of CSREs. (**A**) Number and length of CSREs from each cell type. (**B**) Proportion of genome covered by diverse numbers of CSREs. (**C**) Pairwise overlap of CSREs of all cells and tissues. Orders of rows and columns are consistent with **D**. Group information comes from Roadmap Epigenomics Consortium. (**D**) Hierarchical clustering of cell types or tissues based on pairwise fold enrichment in **C** using Ward linkage. (**E**) Distribution of CSREs on chromosome X across different genders. (**F**) Distribution of CSREs on chromosome X in female ES cells, iPS cells and other groups. Two-sample Wilcoxon tests were used in **E** and **F**.

Twenty-five percent of genome is covered by at least one CSRE, most of which is covered exclusively by only one, demonstrating the cell type or tissue specificity of CSREs (Figure 1B). There are still 5% of genome covered by CSREs from two tissues or cell types and 2% of genome covered by three or more, indicating hierarchical organization of cells or tissues. Overlap of CSREs from common lineages may relate to lineage-sharing functions, whereas co-localization of CSREs from different lineages may imply dynamic behaviors in developmental processes (see subsequent analysis).

To investigate the global view of co-localization between CSREs, we clustered all tissues and cell types by calculating the overlap of CSREs for each pair of tissues or cell types, demonstrating distinct hierarchical organization (Materials and methods, Figure 1C and D). CSREs in a common lineage group tend to be overlapped compared against those between different lineages, implying their common functions within a lineage (Figure 1C). The clustering is consistent to lineage groups, including T cells, muscle, heart, digestive, and brain (Figure 1D and Supplementary Figure S3). It may seem surprising that adult and fetal thymus were clustered together with T cells rather than other tissues. However, it is consistent with the fact that thymus is the place where T cells mature. HeLa cervical carcinoma, A549 lung carcinoma, and HepG2 hepatocellular carcinoma were cluster together, indicating that they share regulatory elements which may relate to cancer functions. While Dnd41 T cell leukaemia and K562 leukeamia were clustered together with blood cells, implying they resemble the original cells rather than other cancer cells. Cell lines from ENCODE such as GM12878, monocytes and HSMM were clustered into corresponding tissues or cell types from Roadmap samples, demonstrating our analysis is robust to data of different projects.

In terms of the distribution of CSREs in all chromosomes, we noticed that chromosome X has the largest coefficient of variation (CV) (Materials and methods, Supplementary Figure S4A). Moreover, in chromosome X, the log_2_(FC) of female samples is significantly higher than that of the male samples (Materials and methods, Figure 1E and Supplementary Figure S4B). Male cells have only one active X chromosome. However, female cells have two copies of X chromosome and one of them is inactivated in somatic cells (Ohno et al., 1959). These inactivation is controlled by epigenomic regulations (Ng et al., 2007). Thus, it is likely to observe more CSREs in chromosome X of female cells. More interestingly, among the female cells, ES cells and iPS cells have CSREs with the highest enrichment on X chromosome, which may relate to the important role of histone modifications in establishment of X inactivation (Barakat and Gribnau, 2010) (Figure 1F).

### CSREs demonstrate distinct functional specificity

We next explored the relationship between CSREs and diverse genomic features. The proportion of CSREs belonging to different genomic features varies across all cell types and tissues (Supplementary Figure S5A). For a vast majority of cell types and tissues, CSREs are enriched in promoters, 5’UTRs, 3’UTRs, exons and introns, and depleted in intergenic regions (Supplementary Figure S5B). As we all know that promoters play key roles in gene regulations. Moreover, UTRs are known to play crucial roles in the post-transcriptional regulation of gene expression (Mignone et al., 2002). Exons and introns can act as splicing regulators (Majewski and Ott, 2002). Enrichment of CSREs in these regions reveals their critical role in regulating corresponding genes with its underlying modification patterns, demonstrating the regulatory potential of CSREs.

CSRE neighboring genes are expected to be cell type-specific ones, and are not likely to be housekeeping ones. As we imagined that, in 79 cell types or tissues, CSRE neighboring genes are significantly depleted in housekeeping genes (FDR < 0.01; Supplementary Figure S6), indicating that genes nearby CSREs tend to perform cell type-specific biological functions. Indeed, in 52 out of 56 cell types or tissues with expression data, CSREs neighboring genes are significantly enriched in cell type-specific ones (FDR < 0.01; Supplementary Figure S7). Moreover, CSRE neighboring genes tend to be enriched in biological functions relating to corresponding cell types or tissues (Figure 2A and Supplementary Table S3). For example, the CSRE neighboring genes of neutrophils and monocytes are significantly enriched in ‘neutrophil activation’; those of T cells tend to be enriched in ‘T cell activation’; those of liver tend to be enriched in ‘organic acid catabolic process’; those of ES and iPS cells tend to be enriched in ‘embryonic organ development’; those of brain tend to be enriched in 'synapse organization'; and those of heart and muscle tend to be enriched in ‘muscle system process’. This analysis highlights the functional specificity of CSREs in regulating expression patterns of genes with cell type-specific functions.

**Figure 2.**
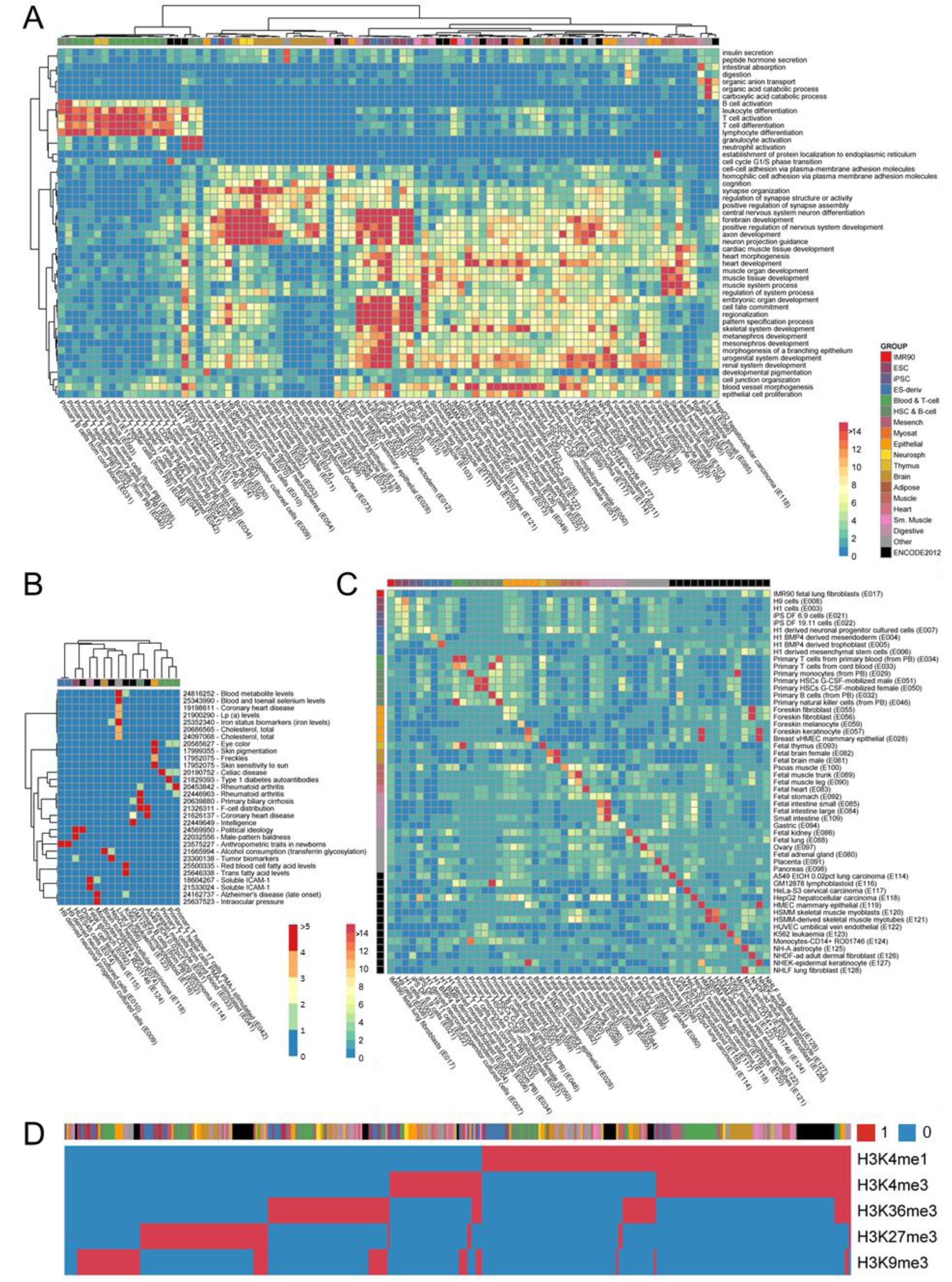
Functional releveance of CSREs with repsect to the enrichment analysis in terms of GO funciton terms, SNP sets associated with phenotypes from GWAS and cell type-specific DNase peaks, and their characterization with diverse histone codes. (**A**) GO enrichment of CSRE neighboring genes and its hierarchical clustering plot. Top 1 enriched GO terms with *q*-value < 1e-6 for each cell type or tissue were selected, and –log_10_(*P*) was used to generate the heatmap. (**B**) Enrichment of GWAS SNP sets of CSREs. Only cases with *q*-value < 0.1 in at least one cell type was shown and –log_10_(*P*) was used to draw the heatmap. (**C**) Overlap between cell type-specific DNase peaks and CSREs. Each row corresponds to cell type-specific DNase peaks, whereas each column relates to CSREs of a cell type or tissue. Color bar was set based on fold enrichment. (**D**) Clustering of binarized centers of all CSRE groups. Groups in **A**, **B**, **C** and **D** are consistent with those in **Figure 1**.

Genome-wide association studies (GWAS) have generated a number of links between single nucleotide polymorphisms (SNPs) and various diseases and traits. However, the mechanism behind such links is still vague. Previous studies have showed that disease-associated variants are relevant to histone modifications, which give clues to the regulatory mechanisms (Trynka et al., 2013). CSREs of a cell type or tissue are defined by five histone modifications with differential signals compared to the remaining ones, suggesting that CSREs may relate to SNPs associated with diseases which are relevant to corresponding cell types or tissues. Interestingly, 30 groups of SNPs are significantly enriched in CSREs of at least one cell type or tissue with false discovery rate (FDR) < 0.1 (Figure 2B and Supplementary Table S4). Moreover, their associated diseases or traits are very consistent with their cellular contexts of CSREs. For example, the SNPs of ‘celiac disease’, ‘type 1 diabetes autoantibodies’ and ‘rheumatoid arthritis’ are significantly located in CSREs of T cells (Cope et al., 2007; Roep and Peakman, 2012; Mazzarella, 2015); the SNPs of ‘primary biliary cirrhosis’ and ‘rheumatoid arthritis’ are significantly enriched in CSREs of B cells (Marston et al., 2010; Zhang et al., 2014); the SNPs of ‘tumor biomarkers’ are significantly enriched in HepG2 hepatocellular carcinoma cell line; the SNPs of ‘skin pigmentation’ and ‘skin sensitivity to sun’ are significantly located in CSREs of foreskin melanocyte primary cells; and the SNPS of ‘cholesterol, total’ and ‘Lp(a) levels’ are significantly enriched in liver. More interestingly, the SNPs of ‘coronary heart disease’ are significantly enriched in CSREs of lung cancer, indicating their potential implications such as the common connection with smoking cessation (Ockene et al., 1990). The SNPs of ‘coronary heart disease’ is also significantly enriched in liver, which is unexpected but consistent with a previous study (Naschitz et al., 2000). While the SNPs of ‘Alzheimer's disease (late onset)’ are enriched in CSREs of monocytes CD14+ rather than those of brain cells, which is also in line with a previous study (Theriault et al., 2015). These results illustrate CSREs can capture the disease-associated variants in corresponding cell types or tissues, which will be valuable for interpreting genetic changes associated with complex diseases.

DNase I hypersensitive sites are comprehensively employed to identify regulatory elements in diverse cell types (Boyle et al., 2008). What’s more, differential DNase-seq footprinting identifies cell type determining transcription factors (Piper et al., 2015). Surprisingly, a majority (69.8%) of CSREs have the highest enrichment in its corresponding cell type-specific DNase peaks, suggesting CSREs could play a crucial role in cell type-specific regulatory activities (Figure 2C). Moreover, ES cells and iPS cells have moderate enrichment, implying their CSREs consist of more repressed regions than those of other cell types.

H3K27ac is a characteristic mark for active enhancers (Creyghton et al., 2010). As H3K27ac tracks were available for only 98 (77%) cell types or tissues, they were not included in our computational pipeline. However, we still found strong enrichment of CSREs in cell type-specific H3K27ac peaks. Specifically, for 78.6% of these 98 cell types or tissues, CSREs are mostly enriched in cell type-specific H3K27ac peaks of their corresponding cell types and tissues than those of others, implying CSREs may behave like active enhancers to regulate target genes (Supplementary Figure S8). ES cells and iPS cells have lower enrichment, consistent with their enrichment with cell type-specific DNase peaks, again suggesting proportion of active regions in their CSREs is lower than those of others.

### Clustering of CSREs of a cell type or tissue based on average specificity signals reveals distinct histone codes

In light of the definition of CSREs, they must be characterized by the specific modification signals in a cell type or tissue. Based on the average specificity scores of each mark for a CSRE, we can cluster the CSREs of a cell or tissue into several groups (Materials and methods). Eighty seven of 127 cell types or tissues harbor three or more groups, indicating CSREs are characterized by different histone codes. To investigate these patterns, we took the center of each group and binarize it to reveal its key marks (Figure 2D). Most (61%) of the groups are contributed by one histone mark, and 34% are associated with two histone marks, demonstrating the distinct histone mark combinations of CSREs.

To better understand the diversity of CSREs, we next studied the groups for specified cell types or tissues. For example, in H1 ES cells, 1641 CSREs were clustered into five groups with distinct histone modification code. Group 1, 2, 4, 5 are characterized by one mark (H3K27me3, H3K4me3, H3K9me3 and H3K4me1), respectively, whereas group 3 involves a combination of two marks (H3K36me3 and H3K9me3) (Figure 3A). The genes closest to CSREs of each group tend to be mutually exclusive, implying most of them are regulated by one distinct histone modification code (Figure 3E). The biggest two clusters are group 4 and 1, covering 27.3% and 25.2% CSREs of H1 cells, respectively. Group 1 harbors extremely high H3K27me3, whereas group 4 owns distinct high H3K9me3, indicating both of them are specific repressed regions (Figure 3B). Cross-sample specificity scores (z-scores) of expression for CSRE neighboring genes of the two groups are indeed significantly lower than those of non-CSRE neighboring genes (*P* = 1.2e-5 and 3.6e-3, respectively, two-sample Wilcoxon test), which is very consistent with their specificity signals from histone modifications (Figure 3F). However, the biological functions of them are quite different. Group 1 is enriched in chromatin states of bivalent/poised TSS, flanking bivalent TSS/enhancer, bivalent enhancer and repressed Polycomb, first three of which are marker states of H1 ES cells (Figure 3D) (Ernst et al., 2011). It is also enriched in DNase peaks, indicating its regulatory potential (Figure 3C). SUZ12 and EZH2 are components of Polycomb Repressive Complex 2 (PCR2), which trimethylates histone H3 on lysine 27 (Margueron and Reinberg, 2011). Strong enrichment of them confirms the specificity of H3K27me3 in group 1. In contrast, CSREs of group 4 are not enriched in peaks of DNase or transcription factors, suggesting it has low regulatory potential. Exclusive enrichment of Heterochromatin state is also consistent with its modification signals. The other two active groups are group 2 and 5. Both of them are enriched in peaks of DNase and H3K27ac, implying they are indeed active regulatory elements. Their specificity scores of expressions of CSRE neighboring genes are both significantly higher than those of non-CSRE neighboring genes (*P* < 2.2e-16 and 2e-11, respectively, two-sample Wilcoxon test). However, CSREs of group 2 is mostly enriched in active TSS state, transcription flanking state, peaks of Pol2, CHD1 and significantly enriched in a large majority of peaks of other transcription factors, indicating its strong relevance to promoter regions. While CSREs of group 5 is significantly enriched in enhancer state, H3K27ac peaks and P300 peaks but not Pol2 peaks, implying its tendency to enhancer nature. The specificity signals and original signals are consistent for group 1, 2, 4 and 5, except group 3 (Supplementary Figure S9). Originally, the median signal of H3K36me3 for both group 3 and 4 do not pass the *P* value threshold 0.01. Thus, only looking at original signals cannot distinguish them well. In contrast, specificity signal of H3K36me3 is higher than those of H3K9me3 in group 3, contradicting group 4 obviously. Consistently, group 3 are significantly enriched in the chromatin state associated with zinc finger protein genes, indicating it does not merely locate in heterochromatin. The specificity scores of expressions of group 3 are also significantly higher than those of group 4 (*P* = 3.7e-12, two-sample Wilcoxon test). All these observations indicate that the distinct characteristics of group 3 against group 4 can be better captured by specificity signals than the original ones.

**Figure 3.**
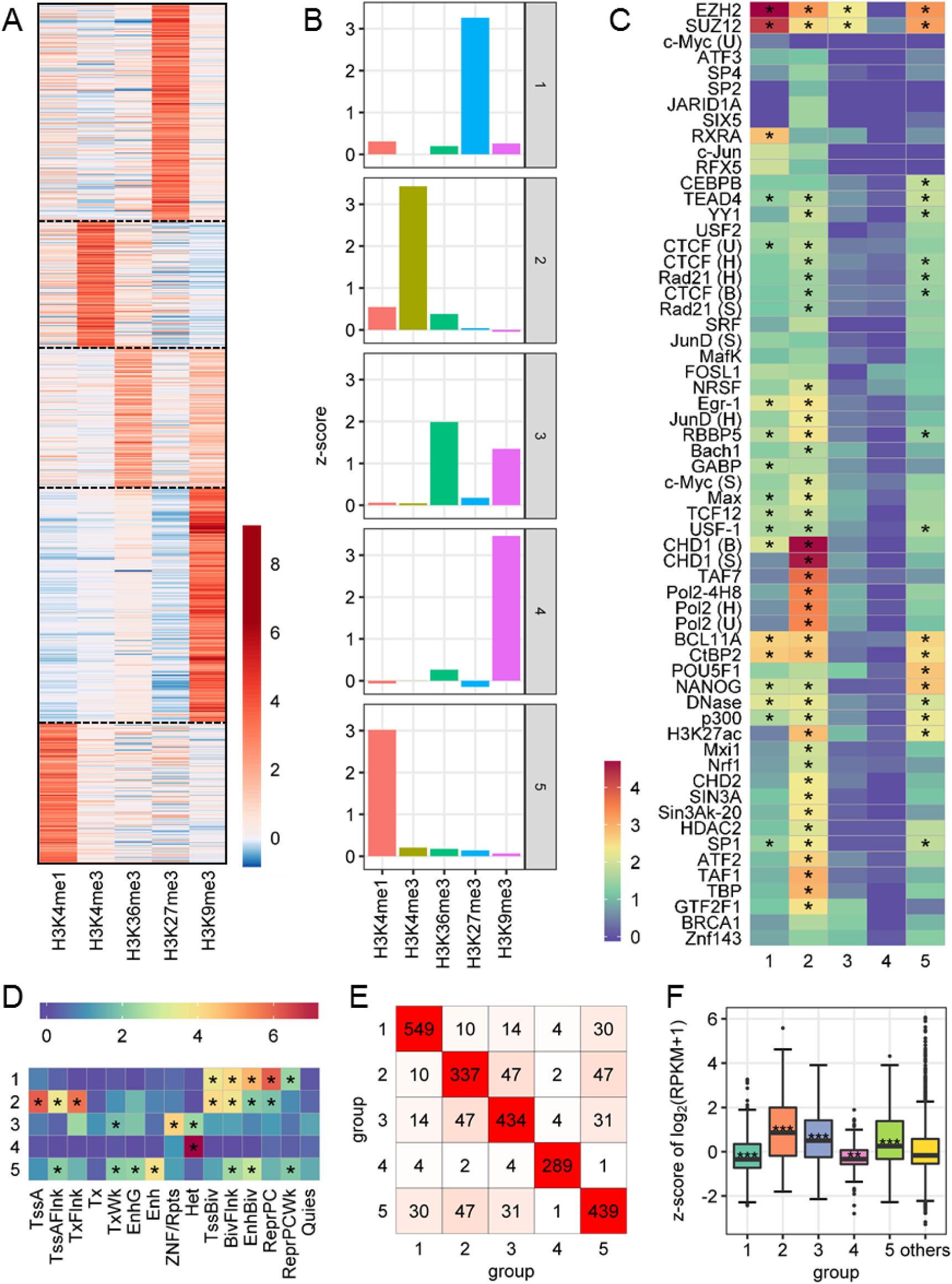
Diverse characteristics of CSRE groups in H1 cells. (**A**) Clustering of CSREs based on their specificity signals. (**B**) Center signals of each group. (**C**) Enrichment analysis for ChIP-seq peaks of transcription factors (TFs) from ENCODE in each group. Letters in brackets (B: Broad Institute; H: HudsonAlpha; S: Stanford; U: University of Texas at Arlington.) were used to distinguish the TF experiments. (**D**) Enrichment analysis of chromatin states in each group. For **C** and **D**, color bar was set based on log_2_(odds ratio + 1) and ‘*’ indicates adjusted *P* < 0.001. Full names of chromatin states are in Supplementary Table S5. (**E**) Pairwise overlap of CSRE neighboring genes from five groups. (**F**) Experssion distribtuion based on z-scores of log_2_(RPKM+1) across cell types and tissues for CSRE neighboring genes. Each group was compared to ‘others’ using two-sample Wilcoxon test. '**' and '***' indicate 0.001 < P < 0.01 and P < 0.001, respectively.

In female fetal brain tissue, the total 3112 CSREs are clustered into four groups, each of them is characterized by one histone mark (H3K4me1, H3K4me3, H3K27me3 and H3K36me3), respectively, which is consistent with the enrichment of chromatin states (Figure 4A, 4B and 4C). The CSRE neighboring genes from group 1, 2 and 4 are distinctly high-expressed, whereas genes around CSREs from group 3 are significantly low-expressed (Figure 4E), which is in line with specificity signals of CSREs. Compared to H1 cells, the fetal brain tissue has substantially more H3K4me3-associated CSREs (59% versus 15%). Another difference of them is that CSREs of group 1, 2 and 4 shared moderate number of genes, which is different from the mutually exclusive characteristic in H1 cells (Figure 4D), implying functional complexity of different cell types or tissues. Interestingly, the CSRE neighboring genes of group 1 and 2 are significantly enriched in brain-associated developmental process, indicating genes regulated by different patterns are likely to work in a cooperative fashion to carry out biological functions in this tissue (Figure 4F). The CSRE neighboring genes of group 3 are enriched in kidney development and cell fate commitment related function terms, demonstrating that brain cell may employ diverse histone marks to regulate various biological processes. All these analysis highlights the diverse functional characteristics of CSREs within a cell type or tissue.

**Figure 4.**
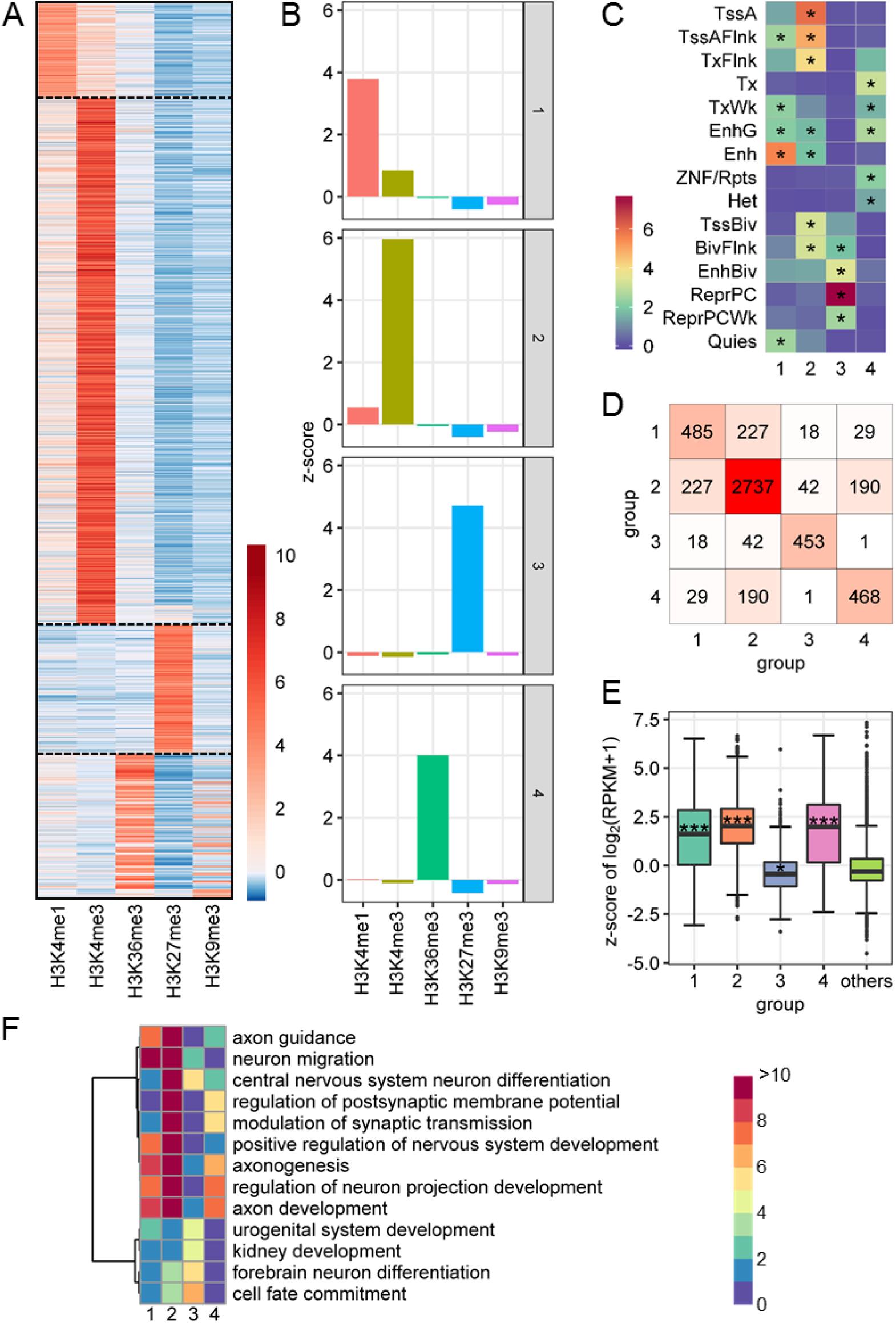
Diverse characteristics of CSRE groups in female fetal brain. (**A**) Clustering of CSREs based on their specificity signals. (**B**) Center signals of each group. (**C**) Enrichment analysis of chromatin states in the four groups respectively. Color bar was set based on log_2_(odds ratio + 1) and ‘*’ indicates adjusted *P* < 0.001. Full names of chromatin states are in Supplementary Table S5. (**D**) Pairwise overlap of CSRE neighboring genes from four groups. (**E**) Expression distribution based on z-scores of log_2_(RPKM+1) across cell types for CSRE neighboring genes. Each group was compared to ‘others’ using two-sample Wilcoxon test. ‘*’ and ‘***’ indicate 0.01 < *P* < 0.05 and *P* < 0.001, respectively. (**F**) Diverse biological processes enriched for the CSRE neighboring genes of the four groups. Top 5 enriched GO terms with q-value < 1e-2 for each group were selected and -log_10_(*P*) was used to generate the heatmap.

### Dynamics of CSREs decipher developmental processes such as normal tissue development and cancer occurrence

In light of the functional specificity, the rare co-localization of CSREs from different lineages with different modification signals can reveal clues for cell or tissue development. For example, we checked the overlapped CSREs from group 1 of H1 cells (potential Polycomb-repressed poised regulators) and group 2 of fetal brain tissue (potential active promoters), and found that the longest one is a 3275bp region (chr22: 28,191,726-28,195,000) located in the first exon and intron of gene MN1 (Figure 5A). Surprisingly, this region is covered by CSREs from iPS cells, H9 derived neuronal progenitor cultured cells, H9 derived neuronal cultured cells, brain germinal matrix and ganglion eminence derived primary cultured neurospheres, as well as some other tissues or cell types, indicating the epigenomic patterns in this region are highly dynamic among diverse cell types. We further inspected the original signals along this region, and found high H3K27me3 signal in H1 cells and high H3K4me3 signal in fetal brain, which is consistent with our specificity signals (Figure 5C). To explore the dynamics of CSREs along the brain developmental process, we further looked into the specificity signals of neuronal progenitor cells and brain germinal matrix. Interestingly, the underlying chromatin modification of CSREs behaves consistently with brain development. In embryonic cells, it is covered by distinctly high H3K27me3 signal, which may repress the transcription of MN1. When the embryonic cells differentiated into neuronal progenitor cells, H3K27me3 are removed, but H3K36me3 and H3K4me3 are established, which may facilitate its transcription. In fetal brain and brain germinal matrix, H3K4me3 is enhanced, which may maintain the expression of MN1. The expression change of MN1 from H1 cells to fetal brain and brain germinal matrix confirms our illustration (Figure 5B). As this region is highly dynamic during brain development, dysregulation of it may relate to abnormal functions in brain. Interestingly, MN1 was indeed reported to be involved in meningioma (Sturm et al., 2016). This example showed the power of utilizing CSREs to explore the dynamic regions between related cell types or tissues.

**Figure 5.**
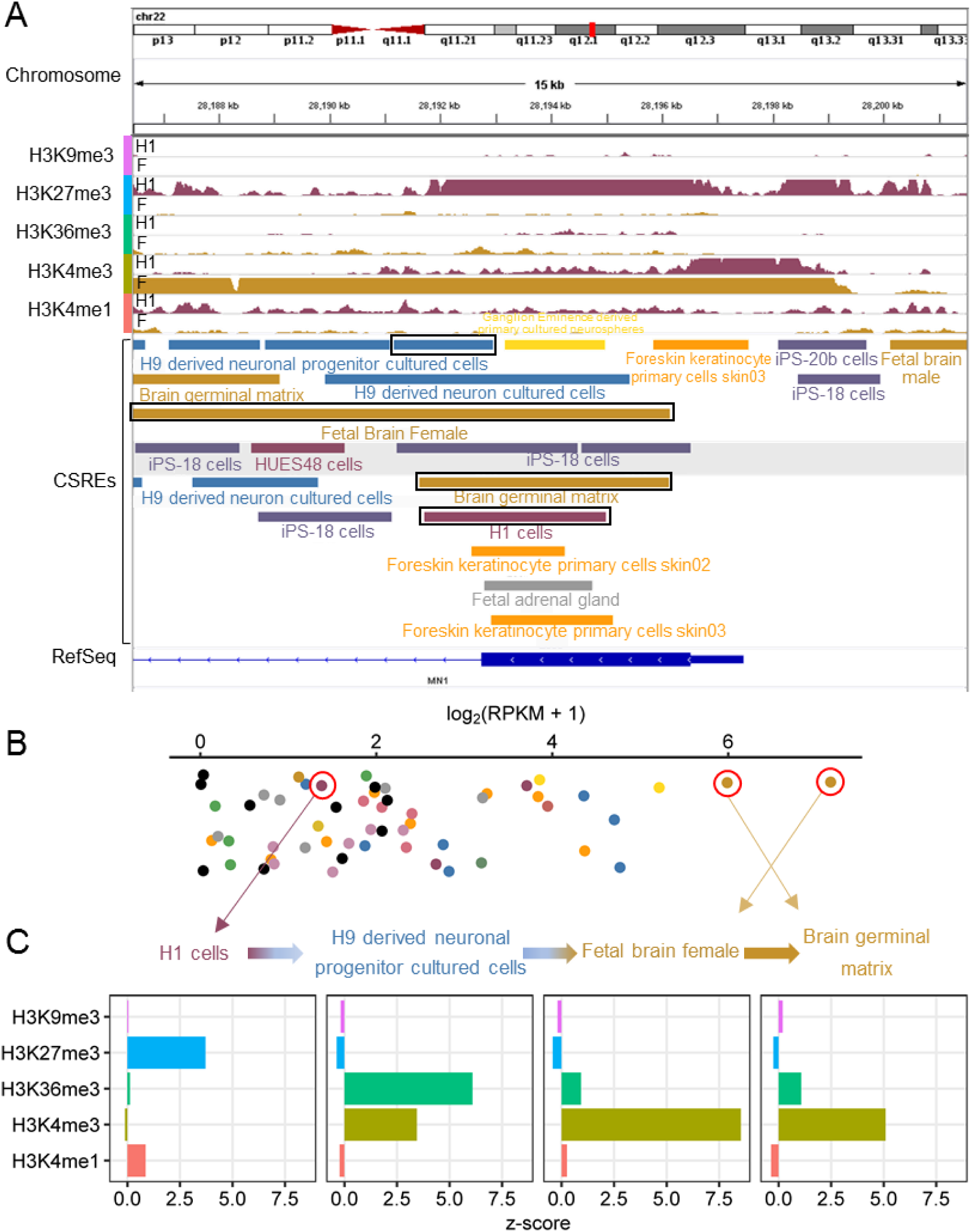
Co-localization of CSREs reveals chromatin dynamics of MN1 during brain development. **(A)** CSREs overlapping the first exon of MN1. The original -log_10_(*P*) tracks of five marks were shown. Extremely high signals were truncated to 40 for visualization. CSREs of cell types were drew in the same colors as those in **Figure 1**. H1: H1 cells; F: Fetal brain female. **(B)** Expression distribution of MN1 in cell types and tissues. **(C)** Specificity signals of four CSREs overlapping the first exon of MN1, which were marked in **A** by rectangles.

In another example, the co-localization of CSREs relates to a cancer gene PRAME in K562 leukemia cells (De Carvalho et al., 2011). CSREs from ES cells, iPS cells, G-CSF-mobilized hematopoietic stem cells and K562 cells co-localize in the last exon and 3’UTR region of PRAME (Figure 6A). By utilizing the specificity signals, we found distinct modification patterns in HUES6 cells, hematopoietic stem cells and K562 cells (Figure 6C). In HUES6 cell, H3K27me3 is distinctly high, which may repress the nearby regions. For hematopoietic stem cells, H3K27me3 signal is moderately low, but H3K4me1 signal is distinctly high, which may perform active regulatory functions. However, K562 gained H3K36me3 additionally, which suggests high expression of PRAME. As expected, K562 has a high expression of PRAME (Figure 6B). Interestingly, it also expresses in A549 lung carcinoma and Hela cervical carcinoma, demonstrating its potential role in cancer.

**Figure 6.**
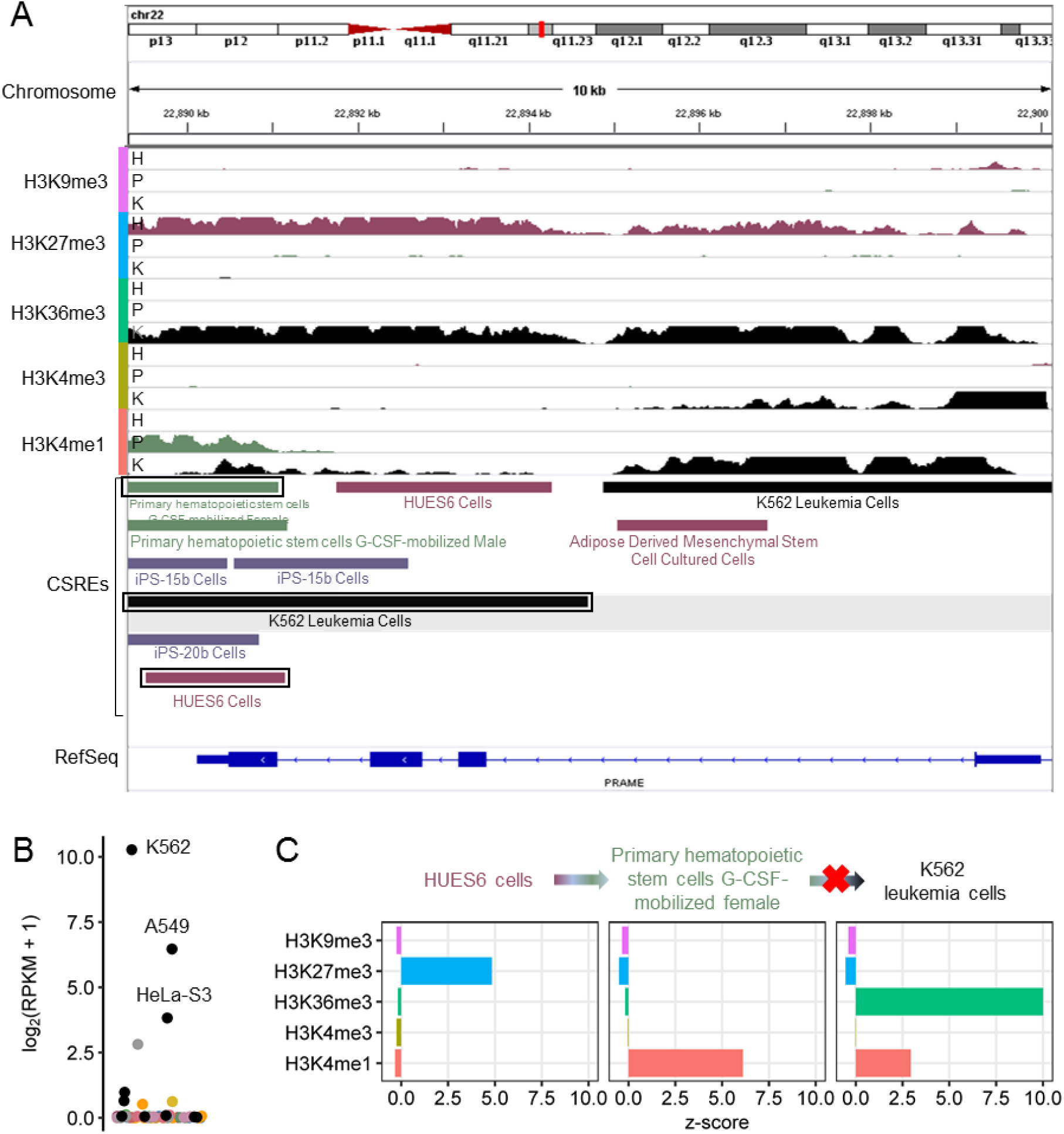
Chromatin dynamics of CSREs relate to cancer gene PRAME in K562 cell line. (**A**) CSREs closing to PRAME. The original -log_10_(*P*) tracks of five marks were shown. Extremely high signals were truncated to 20 for visualization. CSREs of cell types were drew in the same colors as those in **Figure 1**. H: HUES6 cells; P: Primary hematopoietic stem cells G-CSF-mobilized female; K: K562 leukemia cells. (**B**) Expression distribtuion of PRAME. It is highly expressed in the three cancer cell lines. (**C**) Specificity signals of CSREs overlapping the last exon of PRAME, which were marked in **A** by rectangles.

## Conclusion and Discussion

The Human Reference Epigenome Map generated by the Roadmap Epigenomics Consortium provides a unique opportunity to study the regulatory role of chromatin in an integrative way. Genomic regions exhibiting distinctive epigenomic characteristics across different cell types may contribute to cell type-specific gene expression programs. Here we adopted dCMA for addressing continuous signals directly but not binary signals to extract CSREs, which can better distinguish the difference between moderate signals and high signals. We systematically identified cell type-specific regulatory elements (CSREs) for each cell type or tissue of the 127 epigenomes by comparing their five histone modification signals using dCMA. Intriguingly, CSREs relate to known functional genomic features and cell type-specific biological functions, disease-associated SNPs (in the corresponding cell types or tissues), and distinct regulatory potential.

We further characterized CSREs by specificity signals and showed the detailed CSRE groups and diverse histone codes relating to CSREs within each cell type or tissue. CSRE groups show diverse biological relevance and their neighboring genes tend to be regulated by different histone codes, resulting in distinctly high or low expressions. Moreover, the CSRE groups as well as their proportions vary dramatically between different cell types or tissues. The group centers of specificity signals of CSREs captures the main regulatory histone code within a cell type or tissue. Clustering all these group centers of all cells or tissues together gives a systematic view on cell type and tissue-specific histone code.

A vast majority of CSREs locate mutually exclusive with each other, which is consistent with its definition as we expected. The limited overlap of CSREs between cell types or tissues reveals the hierarchical organization underlying cell types and tissues. More interestingly, the rare co-localization of CSREs from different lineages tend to show interesting regions with highly dynamic chromatin events in development of normal tissues or occurrence of disease (e.g., cancer), which are not easy to find with data of small sample size. This observation emphasizes the great value of a comparative method on a large-scale catalogue of epigenomes.

We note that the identification of CSREs relies on the number of cell types or tissues and their common histone mark tracks s used in the pipeline. Larger number of cell types can lead to more reliable CSREs. With epigenomes of more histone marks used, new histone codes relating to CSREs could be identified. However, not all cell types or tissues harbor various histone tracks currently. For the Roadmap reference epigenomes used here, H3K27ac tracks were available for only 77.2% of 127 cell types or tissues. For H3K9ac tracks, the proportion was 48.4%. Moreover, adding both marks would reduce the number of cell types or tissues dramatically from 127 to 49 (Supplementary Table S1). Additionally, H3K27me3 and H3K9me3 antagonize H3K27ac and H3K9ac, respectively. Thus, the extra information of the latter is limited given the former. Hence, we preferred to keeping 127 cell types and tissues with five epigenomes rather than using a relative small number of cell types of tissues with more epigenomes including H3K27ac or H3K9ac. Besides, ChromImpute (Ernst and Kellis, 2015) generated an imputed version of histone tracks for all 127 cell types and tissues. These imputed tracks are generated based on similarity with available tracks and consequently resemble each other more than original tracks, which is contradicted with the definition of CSREs. Hence, we didn’t use the imputed data to detect CSREs. In the future, with the continuous generation of epigenomes of International Human Epigenome Consortium (Stunnenberg et al., 2016), we could get a more accurate and comprehensive map of CSREs on a huge number of cell types and tissues.

## Materials and methods

### Data

We obtained the epigenomic modification tracks (-log_10_(*P* value)) for five histone marks of 127 tissues and cell types (Supplementary Table S1) generated from Roadmap Epigenomics Program (Kundaje et al., 2015) and ENCODE project (Dunham et al., 2012) at http://egg2.wustl.edu/roadmap/data/. These histone marks consist of H3K4me1, H3K4me3, H3K36me3, H3K27me3 and H3K9me3, which relate to regulatory elements, promoters, transcribed chromatin, Polycomb-repressed regions and heterochromatin, respectively. We note that the -log_10_(*P* value) tracks were recommended as the primary signal tracks for analyses by Roadmap Epigenomics Consortium (Kundaje et al., 2015). A series of normalization has been conducted by the consortium to remove bias and make the signal comparable. Reads were mapped to GRCh37/hg19 by PASH (Coarfa et al., 2010) and only uniquely mapping reads were retained. Mapped reads were uniformly truncated to 36bp and filtered by a 36bp mappability track. Corresponding technical/biological replicates were pooled and then uniquely subsampled to a maximum depth of 30 million reads. Finally, MACS2 were used to generate the -log10(*P* value) signal tracks by Poisson distribution with a dynamic parameter. We assumed that the final signals are already comparable for each mark among all cell types as a consequence of all these rigorous preprocessing.

We also downloaded the narrow peaks of those five marks plus H3K27ac and DNase generated by MACSv2.0.10 (Zhang et al., 2008), chromatin states from 15-state models and processed RNA-seq data for available epigenomes at the aforementioned website for validation. We conducted all the following analyses on chromosome 1-22 and X based on assembly GRCh37/hg19.

### Determination of CSREs

We performed the following steps to identify CSREs (Supplementary Figure S1).

#### Step1: Signal correction and z-score transformation

The signals for each track were averaged in non-overlapping 25-bp bins across the whole genome. For each bin on the genome, noise was reduced and z-score was calculated to represent the specificity of each sample compared to the others. Formally, let *v*_*i*,*j*,*k*_ denote the signal value of mark *k* for sample *j* on bin *i*, and *v*_*i*,·,*k*_ denotes (*v*_*i*,1,*k*_, *v*_*i*,2,*k*_, …, *v*_*i*,127,*k*_). To reduce noise on a low signal region, we kept *v*_*i*,·,*k*_ as it is when 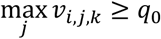. Otherwise, *v*_*i*,·,*k*_ would be set to a vector of all zeros. As -log_10_(*P* value) tracks were used here, we set *q*_0_ to 2, corresponding to a *P* value threshold 0.01. After that, z-score was calculated as 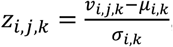, where *μ*_*i,k*_ and *σ*_*i,k*_ are the mean and the standard deviation of *v*_*i*,·,*k*_, respectively. *z*_*i*,*j*,*k*_ carries information about to what extent sample *j* is specific on bin *i* with regard to mark *k*. Note that our method does not need any position-based corrections caused by mappability or GC content. Those corrections are based on an assumption that the ratio of observed signal to real signal is a position-dependent value (Cheung et al., 2011; Benjamini and Speed, 2012). However, as z-score transformation is scale-invariant, it can automatically remove those biases.

#### Step 2: Calculation, normalization and correction of composite score

To integrate information from all marks, we summed up the squares of z-scores on each bin *i* for each sample *j* to get a composite score, which could be represented as 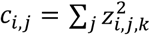. Then we normalized the sum of 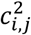 of sample *j* to a constant, say 1000. As the specific sample should not be too many at each bin, the true specific one would have a high composite score which should further be higher than those of other samples. Hence, we corrected *c*_*i,j*_ to 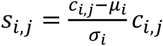, where *μ*_*i*_ and *σ*_*i*_ are the mean and the standard deviation of *c*_*i*,·_, respectively. A high *s*_*i*,*j*_ implies sample *j* is specific on bin *i*.

#### Step 3: Smoothing and CSRE extraction

To borrow information of nearby bins and enhance the signal-to-noise ratio, we adopted wavelet transform to smooth *s*_*i*,*j*_ along the genome. The peaks corresponding to the smoothed *s*_*i*,*j*_ in each cell type were defined as CSREs. Specifically, each CSRE in a cell type is the consecutive bins with length ≥ 1.5kb and *s*_*i*,*j*_ ≥ 0.1. We performed enrichment analysis on CSREs derived from four other sets of parameters to demonstrate their robustness (Supplementary Figure S10, S11 and S12).

### Overlap of CSREs between cell types

To illustrate the spatial relationship of CSREs belonging to different cell types, we investigated the overlap between CSREs. The fold enrichment *e*_*ij*_ of overlap between CSREs of cell type *i* and *j* was defined as

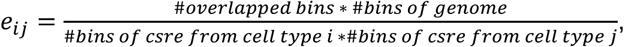

which could be understood as the ratio of observed overlapping bins versus the expected overlapping bins when both bins were sampled randomly along the genome.

We defined a distance *d*_*ij*_ between any two cell types *i* and *j* by *d*_*ij*_ = *M* - *log*2(*e*_*ij*_ + 1), where 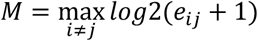.Based on *d*_*ij*_, the cell types were clustered by hierarchical clustering with Ward linkage. We took the *log*2(*e*_*ij*_ + 1) to draw a heatmap, with order of rows and columns consistent to the result of hierarchical clustering. The circular dendrogram was generated by ggtree (Yu et al., 2017). We cut the dendrogram into 25 groups according to the average silhouette scores and compared them with the 19 groups given by Roadmap Epigenomics Consortium using contingency table.

### Distribution of CSREs on chromosomes

We explored the distribution of CSREs on 23 chromosomes (1-22, X) in each tissue or cell type (Supplementary Figure S3). In brief, we calculated the fold change (FC) of observed length of CSREs in each chromosome against expected by randomly sampling them along the genome, and used coefficient of variation (CV) of FC to quantify the variation of a chromosome across all tissues and cell types.

### Mapping CSREs to various genomic features

We examined the potential functional relevance by mapping them to known genetic features. We leveraged RefSeq to build a TxDb objects in Bioconductor on December 16, 2016 and extracted genetic feature therein (Pruitt et al., 2014; Huber et al., 2015). Each transcript named with a prefix of “NM” by RefSeq were regarded as a gene here. Beyond that, we defined six genomic features: promoter, 5’UTR, 3’UTR, exon, intron, intergenic region. Promoters were defined as regions within 2000bp of a transcription start site (TSS) and intergenic regions were composed of base pairs in none of the other five features. We overlapped each CSRE with the fix features and assigned it into one category it has overlap in a priority: promoter > 5’UTR > 3’UTR > exon > intron > intergenic region. Fold enrichment of CSREs and genomic features was calculated in a similar way as aforementioned, but in a 1bp resolution. CSRE neighboring gene was defined as the one with its TSS closed to the CSRE.

### Housekeeping genes

To elucidate the specificity of CSREs, we tested the enrichment of its neighboring genes with known housekeeping genes. Housekeeping genes were obtained from (Eisenberg and Levanon, 2013), and 3793 were kept after mapping to RefSeq genes.

### Specificity of gene expression

RNA-seq data were available for 56 cell types or tissues. Gene expression was calculated as Reads Per Kilobase Million (RPKM) by Roadmap Epigenomics Consortium. We removed 1298 genes with RPKM < 0.5 among all 56 cell types or tissues and kept the left 18497 for analysis. Then the RPKM was transformed to log_2_(RPKM + 1) to reduce skewness. For each gene, we computed its z-scores of log_2_(RPKM + 1) across cell types or tissues and defined them as specificity scores. High positive (or low negative) specificity score indicates specific high (or low) expression for a gene. Cell type-specific genes were defined as those with absolute z-score greater than 2. Difference of z-scores for groups was tested by two-sample Wilcoxon test.

### GO enrichment analysis

We explored the biological function of CSRE neighboring genes by GO enrichment analysis. Each set of concerned genes were mapped to GO terms by org.Hs.eg.db and GO.db Bioconductor packages. Fisher’s exact test was used to get the *P*-value followed by Benjamini-Hochberg correction for each cell type or tissue. Only GO terms with 5 to 500 genes were kept.

### GWAS SNPs enrichment analysis

To investigate the relevance of CSREs and diseases, we tested the enrichment of disease-associated SNPs for CSREs of each cell type. NHGRI GWAS catalogue were obtained from gwascat Bioconductor package (MacArthur et al., 2017). We treated a pair of disease/trait and PubMed Id as a study and tested the overlap of SNPs locating in CSREs for each cell type with those of each study. Significance of the overlap were obtained by Fisher’s exact test with all SNPs in GWAS catalogue as the background. *P*-values were adjusted using the Benjamini-Hochberg correction for each cell type or tissue.

### Cell type-specific DNase peaks and H3K27ac peaks

DNase peaks were available for 53 and 98 cell types or tissues, respectively. Cell type-specific DNase or H3K27ac peaks of a cell type or tissue were defined as part of original peaks that covered by peaks from no more than 4 other cell types or tissues. To demonstrate specificity and functions of CSREs, we depicted their overlap with cell type-specific DNase and H3K27ac peaks. The fold enrichment of overlap between CSREs and cell type-specific DNase or H3K27ac peaks was calculated in a similar way as aforementioned, but conducted in a 1bp resolution.

### Specificity signals and clustering

For each CSRE, we averaged the z-score on its bins for each mark as the specificity signals. Briefly, for CSRE *c* of cell type *j*, the specificity signals can be represented as 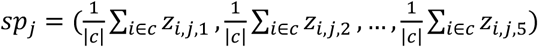.This profile can help to evaluate the specificity of each mark for the CSRE and reveal whether a mark is specific low or high. To explore the diverse subpatterns of CSREs, we performed *k*-means algorithm on their specificity signals. The algorithm was conducted by NbClust R package (Charrad et al., 2014), with distance = “euclidean”, min.nc = 2, max.nc = 10, method = “kmeans” and index = “all”. It used 26 indices and majority rule to determine the best number of *k* and corresponding clusters. The center of each cluster used in the later analysis is based on median rather than mean to get more robust estimation. For each CSRE group, we binarized its center specificity signals with the threshold equal to 1. We drew a heatmap for all CSRE groups in all cell types and tissues based on their binarized centers.

### Overlap analysis with ChIP-seq peaks of transcription factors and chromatin states

We obtained the ChIP-seq peaks of various transcription factors (TF) from ENCODE by AnnotationHub Bioconductor package to inspect their relationship with a set of CSREs. The overlap analysis were conducted by LOLA package (Sheffield and Bock, 2016), with all narrow peaks of the five histone marks (H3K4me1, H3K4me3, H3K36me3, H3K27me3 and H3K9me3) in the corresponding cell type or tissue as the background. LOLA adopted Fisher’s exact test to evaluate the overlapping for each pair of a CSRE set and ChIP-seq peak set. *P*-value were adjusted by Benjamini-Hochberg correction and the odds ratios were used to generate a heatmap. Overlap analysis of chromatin states and a set of CSREs were conducted in the same way.

### Web tool

We built a web tool which is supported by a series of R packages. Users can enter a gene symbol or chromatin region, zoom in/out, and move left/right to jump to their interested regions (Supplementary Figure S13). Users can also select a sub-region and check the corresponding CSREs shown below.

